# Acute inflammation alters energy metabolism in mice and humans: Role in sickness-induced hypoactivity, impaired cognition and delirium

**DOI:** 10.1101/642967

**Authors:** John Kealy, Carol Murray, Eadaoin W. Griffin, Ana Belen Lopez-Rodriguez, Dáire Healy, Lucas Silva Tortorelli, John P. Lowry, Leiv Otto Watne, Colm Cunningham

## Abstract

Systemic infection triggers a spectrum of metabolic and behavioral changes, collectively termed sickness behavior, that while adaptive for the organism, can affect mood and cognition. In vulnerable individuals, acute illness can also produce profound, maladaptive, cognitive dysfunction including delirium, but our understanding of delirium pathophysiology remains limited. Here we used bacterial lipopolysaccharide (LPS) in C57BL/6J mice and acute hip fracture in humans to address whether disrupted energy metabolism contributes to inflammation-induced behavioral and cognitive changes. LPS (250 μg/kg) induced hypoglycemia, which was mimicked by IL-1β (25 μg/kg) but not prevented in IL-1RI^-/-^ mice, nor by IL-1RA (10 mg/kg). LPS suppression of locomotor activity correlated with blood glucose concentration, was mitigated by exogenous glucose (2 g/kg) and was exacerbated by 2-deoxyglucose glycolytic inhibition, which prevented IL-1β synthesis. Using the ME7 model of chronic neurodegeneration, to examine vulnerability of the diseased brain to acute stressors, we showed that LPS (100 μg/kg) produced acute cognitive dysfunction, selectively in those animals. These acute cognitive impairments were mimicked by insulin (11.5 IU/kg) and mitigated by glucose, demonstrating that acutely reduced glucose metabolism impairs cognition in the vulnerable brain. To test whether these acute changes might predict altered carbohydrate metabolism during delirium, we assessed glycolytic metabolite levels in cerebrospinal fluid (CSF) in humans during delirium, triggered by acute inflammatory trauma. Hip fracture patients showed elevated CSF lactate and pyruvate during delirium, consistent with altered brain energy metabolism. Collectively the data suggest that disruption of energy metabolism drives behavioral and cognitive consequences of acute systemic inflammation.

## Introduction

Systemic infection triggers a spectrum of metabolic and behavioral changes, termed sickness behavior, which includes fever, lethargy, hypophagia, anhedonia. Sickness behavior is an evolutionarily conserved response to illness and represents a reprioritization by the organism to conserve energy and maximize the probability of recovery (Dantzer, 2004). Systemic administration of the bacterial endotoxin lipopolysaccharide (LPS) can induce sickness behavior in humans (Schedlowski et al., 2014; Draper et al., 2017) and rodents (Teeling et al., 2007; Carlton and Demas, 2017) and despite not readily crossing the blood-brain barrier (Banks and Robinson, 2010), LPS increases central pro-inflammatory cytokines including interleukin-1β (IL-1β) and tumor necrosis factor alpha (TNF-α) (Skelly et al., 2013), and alters local field potential (Semmler et al., 2008; Mamad et al., 2018). Peripheral inflammatory status is communicated to the brain via: direct vagal signaling to the brainstem and hypothalamus; macrophage activation in the circumventricular organs lacking a patent BBB, leading to secretion of inflammatory mediators into the parenchyma; and activation of endothelial cyclooxygenases to secrete lipophilic prostaglandins directly into the parenchyma (Dantzer, 2018). Manipulation of prostaglandin-dependent mechanisms revealed neuroanatomical pathways underpinning sickness responses (Saper et al., 2012) but the molecular basis for acute LPS-induced suppression of activity is poorly understood.

Sickness behavior sometimes encompasses cognitive impairment: peripheral LPS or IL-1β administration can affect synaptic plasticity and hippocampal-dependent learning and memory (Yirmiya and Goshen, 2011), although the relative preservation of cognitive function is striking given the overt suppression of spontaneous behavior (Cunningham and Sanderson, 2008; Skelly et al., 2018). However, when inflammatory insults are severe, or occur in older age or during evolving dementia, they may induce delirium (Elie et al., 1998). Delirium is an acute onset and fluctuating syndrome characterized by inability to sustain attention, reduced awareness and perception, and profound cognitive impairment (American Psychiatric Association, 2013), affecting approximately 1/5 hospital inpatients (or 1/3 for those >80 years age) (Ryan et al., 2013). Delirium is associated with extended hospitalization, subsequent cognitive decline, and increased risk for dementia but the neurobiological understanding of delirium is limited.

We have modeled delirium, using superimposition of LPS upon models of neurodegeneration (Field et al., 2012; Murray et al., 2012; Lopez-Rodriguez et al., 2018) to produce acute onset, fluctuating deficits in relevant cognitive domains (Davis et al., 2015). These LPS-associated deficits are absent in normal animals but susceptibility to LPS-induced cognitive impairment increases as a function of the underlying neurodegenerative state of the brain (Davis et al., 2015). LPS-induced deficits are prostaglandin-dependent and can be mimicked by systemic administration of IL-1β (Griffin et al., 2013) or TNF-α (Hennessy et al., 2017), and reduced by systemic administration of IL-1 receptor antagonist (IL-1RA) (Skelly et al., 2018) suggesting that IL-1β may affect cognition via a peripheral route. One possibility is that acute sickness impinges on cerebral metabolism through systemic metabolic changes; cerebral glucose uptake is reduced in a rat model of LPS-induced sepsis (Semmler et al., 2008), carbohydrate metabolism is decreased post-LPS (Irahara et al., 2018), and IL-1 has been demonstrated to induce hypoglycemia (Del Rey et al., 2006). Systemic hypoglycemia impacts on central glucose levels (Kealy et al., 2015), which in turn can affect neuronal activity and may be especially detrimental if the brain is already compromised during evolving neurodegenerative pathology.

Therefore, we hypothesized that LPS induced-disturbances in glucose metabolism would drive suppression of activity and cognitive impairment in mice. We assessed locomotor activity and working memory, while manipulating glucose metabolism with LPS, 2-deoxyglucose (2-DG) and insulin to determine effects of altered glycemic status on sickness behavior and cognitive impairments in disease-naïve and ME7 mice. Finally, we analyzed glycolysis products in the cerebrospinal fluid (CSF) of inflammatory trauma (hip fracture) patients to assess brain energy metabolites in humans with inflammation-induced delirium. The findings show that altered glycemic status causes disruption of brain function in mice and that brain carbohydrate metabolism is also disrupted during delirium in patients.

## Materials and Methods

### Animals

Female c57BL/6J and mixed sex IL-1R1^-/-^ mice aged 5-8 months (Harlan, UK) were housed at 21 °C with a 12/12-hour light-dark cycle (lights on 08:00-20:00) with food and water available *ad libitum*. All animal experiments were in accordance with European Commission Directive 2010/63/EU and were performed following ethical approval by the TCD Animal Research Ethics Committee and licensing by the HPRA.

### ME7 prion model of neurodegeneration

Mice were anaesthetized using Avertin (2,2,2-tribromoethanol 50 % w/v in tertiary amyl alcohol, diluted 1:40 in H_2_O; 20 ml/kg, i.p.; Sigma, Ireland) and placed in a stereotaxic frame (David Kopf Instruments, Tujunga, CA, USA). 1 μl of 10% w/v ME7-infected c57BL/6J or 10% w/v normal brain homogenate (NBH) in sterile PBS was infused into the dorsal hippocampus at −2.0 mm (A/P); ±1.6 mm (M/L); −1.7 mm (D/V) from Bregma as described previously (Murray et al., 2011; Murray et al., 2012). Mice recovered in a heated chamber, then returned to their home cage where their drinking water was supplemented with sucrose (5 % w/v) and Carprofen (0.05 % v/v; Rimadyl, Pfizer, Ireland).

### Treatments

Mice were injected intraperitoneally (i.p.) with one or a combination of the following treatments using sterile saline as a vehicle: LPS from *Salmonella equine abortus* (100μg/kg or 250 μg/kg; Sigma, Ireland), IL-1β (25 μg/kg; R&D Systems, Minneapolis, MN, USA), IL-1RA (10 mg/kg; Kineret, Biovitrum, Sweden), glucose (2g/kg; Sigma, Ireland), 2-DG (2 g/kg; Sigma, Ireland), and insulin (11.5 IU/kg (400 μg/kg); Sigma, Ireland). LPS was administered 2 hours prior to open field behavior or 3 hours prior to the T-maze task. Glucose was administered 30 minutes before any behavioral task.

### Behavioral assessment

Spontaneous activity was assessed by observing locomotor activity in an open field as previously described (Murray et al., 2013). Briefly, mice were allowed to freely and individually explore an open field arena (58 × 33 × 19 cm), which was divided into squares (10 × 10 cm). Over the course of 3 minutes, the number of squares crossed by each mouse was counted.

Cognitive performance was assessed using an escape-from-water alternation task in a paddling T-maze as described previously (Murray et al., 2012; Skelly et al., 2018). Briefly, this working memory task involves two runs per trial. On the first run, mice only have one of the T-maze arms available to them, which has an exit at the end of it to escape from the shallow water. On the second run, mice are given a choice between the two arms with the exit now in the opposite arm to the first run. Mice were trained (2 blocks/day, 5 trials/block, 2 runs/trial) until they performed with at ≥80 % success. They were then pharmacologically challenged and tested on the same day of the challenge. There were 3 blocks of 5 trials post-challenge (corresponding to 3-5 (+3), 5-7 (+5), and 7-9 (+7) hours post-challenge) and 2 blocks for insulin, due to its rapid action on blood glucose (corresponding to 40-160 min (+1 h) and 160-300 min (+3 h) post-challenge). All mice underwent recovery testing (2 blocks of 5 trials) on the following day.

### Blood glucose measurements

For serial blood measurements, mice were placed in a plastic restrainer, the tail vein was dilated using warm water and lanced using a 30 G needle. Glucose was measured using a veterinary glucometer (AlphaTRAK 2, Zoetis, USA). Terminal blood glucose measurements were made following sodium pentobarbital overdose and incising the right ventricle, immediately before transcardial perfusion. Blood glucose readings were higher on the veterinary glucometer compared to a clinical glucometer (data not shown) but basal levels were broadly in line with other studies (Del Rey et al., 2006; Del Rey et al., 2016; Chakera et al., 2018; Tooke et al., 2019).

### CSF sampling and analysis

In mice, CSF was collected under terminal anesthesia. Mice were placed in a stereotaxic frame and the cisterna magna accessed by lowering the incisor bar on the animal’s head to angle it downwards at 45° from horizontal. Using a small volume insulin syringe (BD Micro - Fine™ + 0.3ml Insulin Syringe Demi), a freehand puncture was performed slowly to avoid brain stem damage and blood contamination. Approximately 5 μl was collected in 0.5 ml microcentrifuge tubes.

### Hip fracture patient cohort

Hip fracture is a frequent occurrence in frail, elderly populations. Delirium occurs with high prevalence in these patients (Marcantonio, 2017) and since these patients, in many centers, receive spinal anesthesia for hip-fracture repair surgery, this offers an opportunity for CSF collection allowing assessment of the impact of this acute inflammatory trauma on CSF markers of brain energy metabolism in older individuals. CSF was collected from hip fracture patients acutely admitted to Oslo University Hospital after informed consent from the patient and/or proxy (if patients were unable to consent due to cognitive impairment), as approved by the Regional Committee for Medical and Health Research Ethics (South-East Norway; REK 2009/450). The presence of delirium was assessed in all participants using the Confusion Assessment Method (CAM) (Inouye et al., 1990) based on a 10-to 30-minute interview with participants and information from relatives, nurses and hospital records. One geriatrician and one old age psychiatrist independently evaluated whether participants met the ICD-10 criteria for dementia prior to the fracture, based on all available data, as explained earlier (Watne et al., 2014b). CSF was collected in propylene tubes at the onset of spinal anesthesia. Samples were centrifuged, aliquoted and stored at –80° C, as previously described (Watne et al., 2014a).

Samples were defrosted and transferred to CMA Microvials (CMA Microdialysis AB, Sweden). In mice, where sample volumes were too small (<3 μl), samples from the same treatment groups were pooled. Glucose, lactate and pyruvate (the latter in humans only) concentrations were determined using a CMA600 Microdialysis Analyzer (CMA Microdialysis AB, Sweden). Basal levels of CSF glucose were in the range expected from other studies (Horn and Klein, 2010; Nakamura et al., 2017; Tang et al., 2017), as was the case for CSF lactate (Horn and Klein, 2010). The lower limits of detection were: Glucose (0.1 mmol/l); lactate (0.1 mmol/l); and pyruvate (10 μmol/l). Since the ability to detect pyruvate is an indication of its concentration, non-detected samples were included at 0 μmol/l as a conservative measure.

### Statistical analysis

Statistical analysis was performed in GraphPad Prism 5 and IBM SPSS Version 25. Pairs of groups were measured using t-tests and all multiple comparisons were made using ANOVAs, paired and repeated measure variants were used as appropriate. Full statistical analyses and experimental numbers are included in figure legends. Where data were found to violate the assumptions of t-tests, Mann-Whitney U-tests were used instead.

## Results

### LPS and IL-1β both robustly reduce systemic glucose concentrations

The effects of systemic infection on spontaneous activity have largely been attributed to changes in cytokine signaling (Dantzer, 2018). IL-1 is reported to act at the endothelium or at forebrain targets to mediate LPS-induced suppression of motivated behaviors (Liu et al., 2019) but precisely how suppression of exploratory activity occurs is unclear. Here, we examined LPS-induced suppression of activity (see figure 1A) and confirmed that LPS (250 μg/kg; i.p.) significantly increased plasma IL-1β levels at 2 and 6 hours post-challenge in c57BL6/J mice compared to saline-treated controls (Figure 1B) and also reduced blood glucose levels by >50 % by 7 hours post-challenge (Figure 1C). Blood glucose was decreased as early as 3 hours following LPS and had not fully returned to baseline levels by 24 hours. This reduction is not explained by suppression of feeding since blood glucose declines rapidly here (Figures 1B and E) but only begins to decrease after 6-12 hours of fasting in healthy c57BL6/J mice (Champy et al., 2004). We investigated the contribution of IL-1β signaling to LPS-induced hypoglycemia. IL-1β (25 μg/kg; i.p.) reduced blood glucose, with a more rapid induction and earlier nadir than LPS (Figure 1D). IL-1β-induced reductions in glucose were completely blocked by IL-1RA (10 mg/kg i.p.). Therefore IL-1β is sufficient to reduce systemic glucose levels.

**Figure 1.**
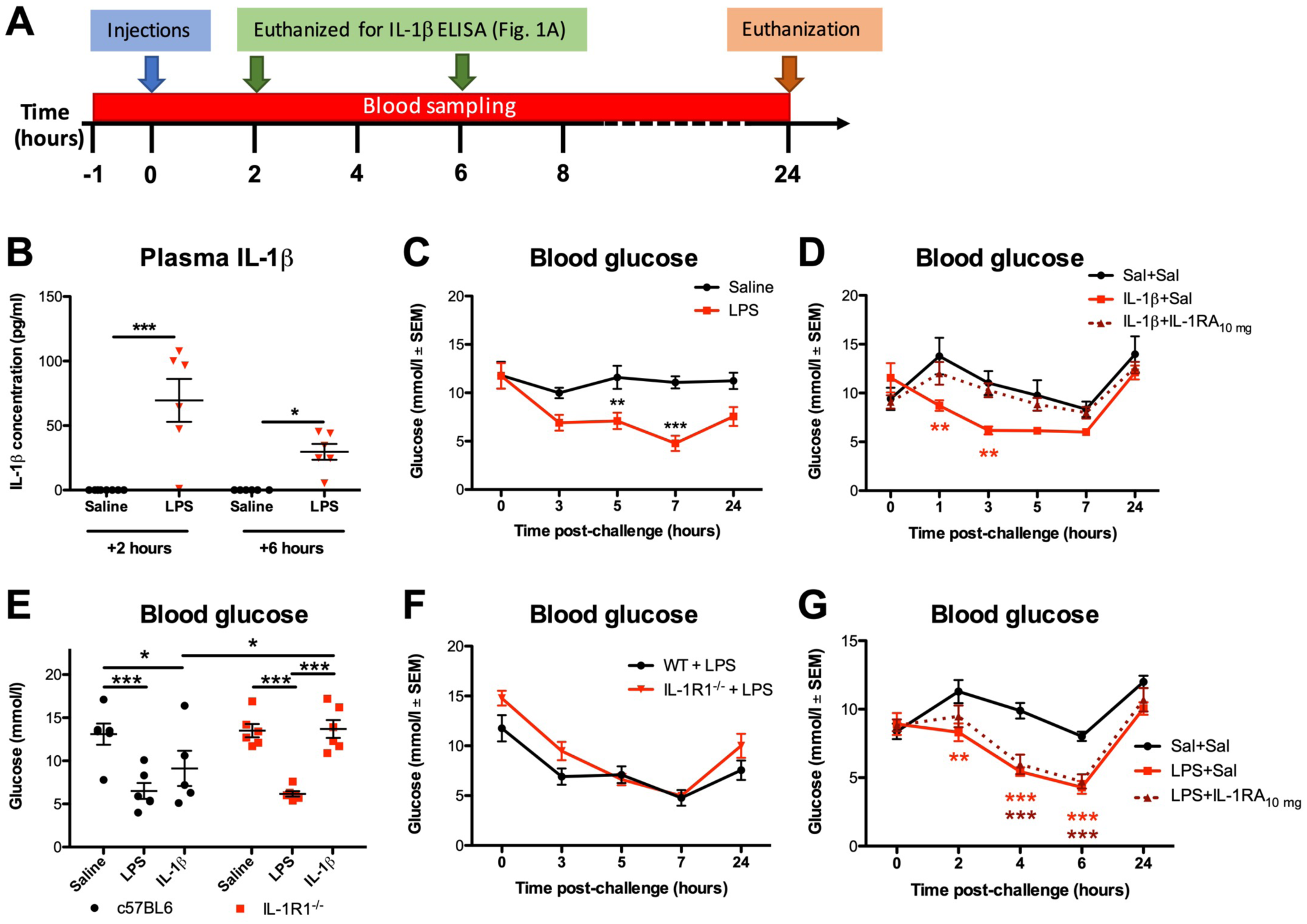
LPS and IL-1β significantly lower blood glucose concentrations. **A:** Timeline for treatments and sampling times. Blood sampling was from tail vein, aside from the 24 hour time point where glucose levels were measured from right atrial blood before transcardial perfusion. In one cohort, mice were euthanized at 2 and 6 hours post-LPS challenge to collect plasma for the IL-1β ELISA. **B:** LPS treatment (250 μg/kg, i.p.) significantly increased plasma IL-1β (*F*_*(1,22)*_ = 36.71; *p* < 0.0001; n = 8 for saline/2 h group; n = 6 for other groups). **C:** LPS treatment (n = 7) significantly reduced glucose levels over 24 h compared to saline controls (0.9 %, i.p.; n = 6); main effect of treatment (*F*_*(1,44)*_ = 24.10; *p* = 0.0005). **D:** IL-1β (25 μg/kg, i.p.; n = 7) reduced systemic glucose and IL-1β’s effect can be blocked using IL-1 receptor antagonist (IL-1RA; 10 mg/kg, i.p.; n = 7). Main effect of treatment (*F*_*(2,85)*_ = 3.843; *p* = 0.0420) and ** denotes significantly lower glucose levels in IL-1β+Saline-treated mice compared to controls (n = 6) at 1 and 3 h post-challenge. **E:** c57BL6/J mice, both LPS (n = 6) and IL-1β (n = 5) significantly reduced blood glucose 4 h post-challenge versus saline controls (n = 6) while in IL-1R1^-/-^ mice, LPS (n = 6) but not IL-1 (n = 6) significantly reduced blood glucose versus controls (n = 6). Significant pairwise comparisons by Bonferroni *post hoc* test after a main effect of treatment (*F*_*(2,29)*_ = 21.81; *p* < 0.0001) are annotated by * (p < 0.05) and *** (p < 0.001). **F:** Time course of changes in blood glucose in IL-1R1^-/-^ (n = 5) and c57BL/6J mice (n=7; significant effect of genotype, *F*_*(1,40)*_ = 5.673; *p* = 0.0385, but no pairwise differences at any time point). **G:** IL-1RA (10 mg/kg, n=12) administered immediately after LPS treatment modestly attenuated LPS-induced reductions in glucose (*F*_*(2,132)*_ = 16.18; *p* < 0.0001) but this was a transient effect (*F*_*(4,132)*_ = 39.08; *p* < 0.001). There was a significant interaction of treatment and time (*F*_*(8,132)*_ = 3.502; *p* = 0.0011) and *post hoc* tests indicated that LPS+Saline-treated mice (n = 12) had significantly lower blood glucose levels versus saline (n = 12) at 2, 4 and 6 h post-challenge while LPS+IL-1RA (n = 12) did not significantly decrease glucose levels compared to controls until 4 h. All annotated Bonferroni *post hoc* tests were performed after significant main effects or interactions in ANOVA analysis: \**p* < 0.05; \*\**p* < 0.01; \*\*\**p* < 0.001.

To test whether IL-1β is necessary for LPS-induced hypoglycemia, we administered LPS (250 μg/kg; i.p.) and IL-1β (25 μg/kg; i.p.) to IL-1 receptor-1 knockout (IL-1R1^-/-^) mice and c57BL6/J wild type (WT) controls. Blood glucose measurements were taken 4 hours post-challenge, to ensure robustly decreased glucose (Figure 1C; 1D). LPS and IL-1β again reduced blood glucose in WT mice. Although IL-1β-induced hypoglycemia was prevented in IL-1R1^-/-^ mice, LPS-induced reductions in glucose were statistically indistinguishable from those in WTs (Figure 1E). Moreover, the time course of LPS-induced glucose reduction was highly overlapping in WT and IL-1RI^-/-^ mice (Figure 1F). IL-1β antagonism with IL-1RA has been reported to attenuate LPS-induced hypoglycemia (Del Rey et al., 2006). Here, IL-1RA (10 mg/kg) showed only a very modest and temporary protective effect against 250 μg/kg LPS-induced decreases in glucose 2 hours post-challenge and no effect thereafter, as detailed in Figure 1G and associated legend. Therefore systemic IL-1β is sufficient to lower blood glucose but it is not indispensable for LPS-induced decreases in glucose.

### Blood glucose concentration is a major determinant of LPS-induced acute hypoactivity

IL-1β has been reported as the major driver of sickness behavior (Matsuwaki et al., 2017; Dantzer, 2018). We sought to understand whether IL-1β signaling or decreases in glucose might be the proximate cause of LPS-induced hypoactivity. LPS-induced hypoactivity in c57BL6/J mice (squares crossed/3 minutes) was significantly positively correlated with blood glucose levels (Figure 2A), with low blood glucose predicting low activity in LPS-treated mice (r^2^=0.4824, p=0.002), while no correlation was present between blood glucose and activity in saline-treated animals. In a separate experiment, LPS-treated mice showed significantly less spontaneous activity in the open field compared to saline-treated controls but when inactive mice were prompted to move, using a gentle finger nudge, similar levels of activity were observed in both groups (Figure 2B) demonstrating that these animals are not unable to move or prevented from moving due to reduced energy availability. Rather, LPS-induced hypoactivity (Figure 2A) reflects reduced spontaneous activity.

**Figure 2:**
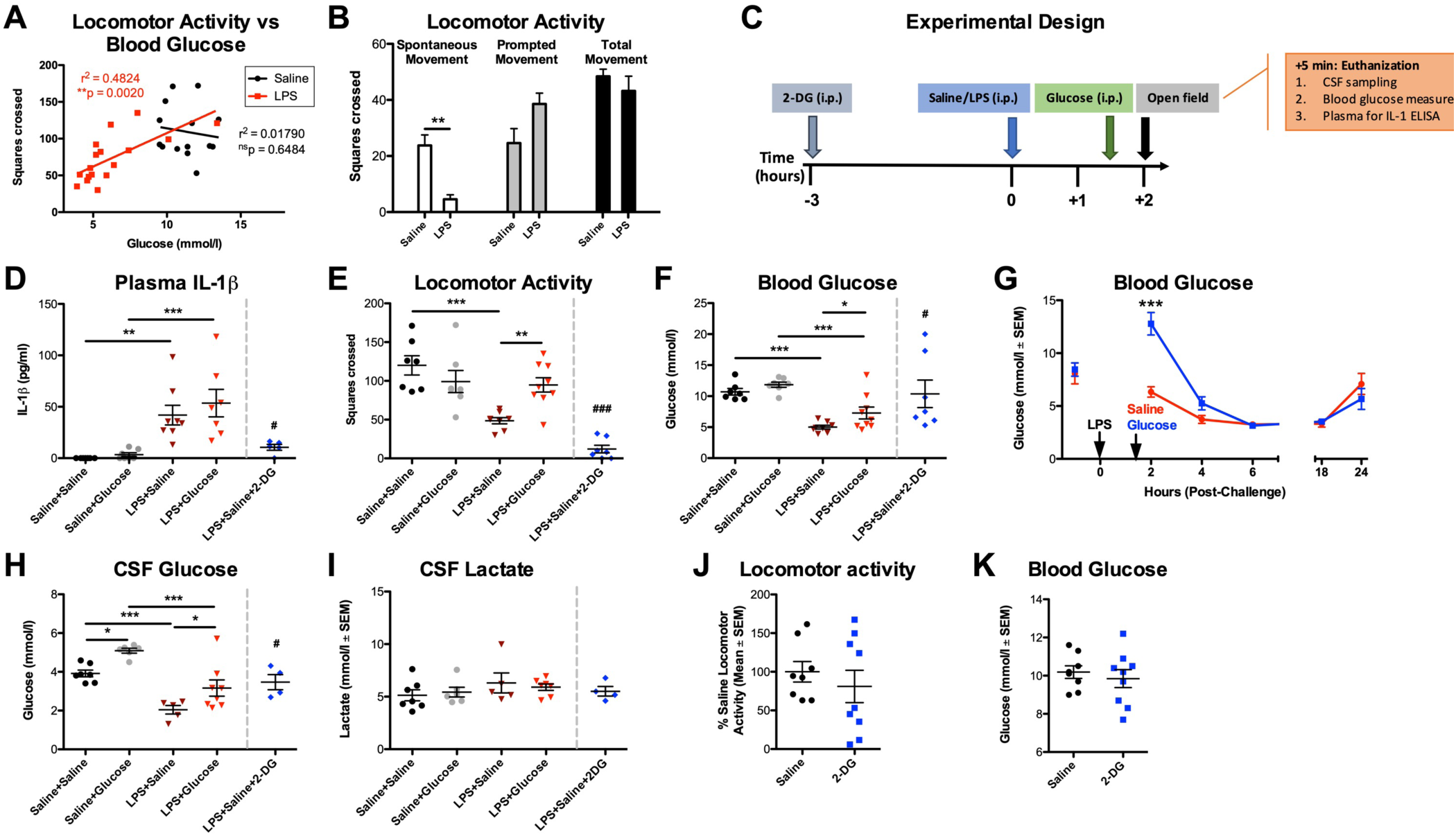
Low blood glucose concentration drives LPS-induced hypoactivity. **A:** Linear regression analyses of locomotor activity (squares crossed/3 min) versus blood glucose concentration (mmol/l) in animals challenged with saline (n = 14) and LPS (n = 17). Blood glucose concentrations significantly correlated with locomotor activity in LPS-treated mice. **B:** LPS significantly reduces spontaneous activity in the open field compared to saline-treated controls. Prompting inactive mice to move by gently nudging them with a fingertip results in similar levels of activity, showing that LPS mice are capable of moving but are not motivated to do so. **C:** Timeline for treatments and sampling times. Glucose (2 g/kg; i.p.) was administered 1.5 hours post-LPS challenge (250 μg/kg; i.p.) and open field behavior was measured 2 hours post-LPS challenge. 5 minutes after open field testing, mice were euthanized, CSF samples taken, blood glucose levels assessed, and plasma collected for IL-1β ELISA. In one group, 2-deoxyglucose (2-DG; 2 g/kg; i.p.) was given 3 hours prior to LPS. **D:** LPS (250 μg/kg, i.p.; n = 8) induced IL-1β production (*F*_*(1,25)*_ = 29.88; *p* < 0.001), which was unaffected by glucose co-administration (n = 7; 90 minutes post-LPS) but blocked by 2-DG administration (i.p., n = 5, ^#^*p* = 0.0296 vs. LPS+Saline). **E:** Locomotor activity was suppressed by LPS (main effect of LPS: *F*_*(1,27)*_ = 13.39; *p* = 0.0011) but rescued by glucose co-administration (interaction between treatments: *F*_*(1,27)*_ = 10.48; *p* = 0.0032). ** denotes significant difference between LPS+Glucose (n = 9) and LPS+Saline (n = 8) and these were not significantly different to Saline+Saline (n = 7) or Saline+Glucose controls (n = 7). 2-DG+LPS completely suppressed locomotor activity in LPS-treated mice (*t*_*(13)*_ = 5.766; ^###^*p* < 0.0001 vs. LPS+Saline). **F:** Blood glucose was suppressed by LPS (main effect: *F*_*(1,27)*_ = 60.00; *p* < 0.0001) and modestly increased by glucose (main effect: *F*_*(1,27)*_ = 6.721; *p* = 0.0152) and *post hoc* tests showed that LPS+Glucose was significantly different to LPS+Saline. **G:** Glucose treatment 1.5 h after LPS provided significant but transient protection against LPS-induced hypoglycemia. **H:** CSF (from the same animals) showed a main effect of LPS (*F*_*(1,22)*_ = 39.85; *p* < 0.0001) and a strong main effect of glucose (*F*_*(1,22)*_ = 14.57; *p* = 0.0009). LPS+Glucose was significantly different to LPS+saline in *post hoc* analysis. **I:** CSF lactate levels (same animals), were not altered by the treatments described. In LPS-naïve mice, 2-DG on its own does not significantly affect **J:** spontaneous activity nor does it: have any effect on **K:** blood glucose. Significance levels for Bonferroni *post hoc* tests: \**p* < 0.05; \*\**p* < 0.01; \*\*\**p* < 0.001.

We, then hypothesized that LPS-induced hypoactivity would be mitigated by treatment with glucose (2 g/kg; i.p.). In our experimental design (Figure 2C), we also included a separate group of mice that received LPS+2-deoxyglucose (2-DG), which inhibits glucose-6-phosphate isomerase to prevent glycolysis, which in turn blocks macrophage synthesis of IL-1β (Tannahill et al., 2013). LPS robustly produced IL-1β, reduced blood and CSF glucose levels, and suppressed activity. Glucose treatment had no effect on IL-1β production (Figure 2D) but significantly improved locomotor activity (Figure 2E) and increased circulating glucose concentration (Figure 2F). Administering glucose to mice, following LPS treatment, only transiently protected against LPS-induced decreases in blood glucose levels (Figure 2G) as has previously been shown (Del Rey et al., 2006). In CSF, LPS significantly reduced glucose concentrations and, similar to the periphery, glucose treatment significantly protected against this (Figure 2H). However, CSF lactate remained the same in all groups (Figure 2I). Therefore, LPS reduces CSF glucose concentration by approximately 50 % and this can be mitigated by systemic glucose administration, with concomitant rescue of spontaneous activity. As predicted, 2-DG blocked LPS-induced IL-1β secretion (Fig. 2D) and yet hypoactivity remained striking (Fig. 2E). Therefore, animals with high IL-1β can remain spontaneously active if glucose concentration is temporarily boosted and, despite LPS producing no IL-1β, animals additionally exposed to an inhibitor of glycolysis show little or no locomotor activity. These findings were replicated in a separate cohort of mice using a separate batch of LPS (100 μg/kg; i.p.; data not shown). While 2-DG had a significant effect on spontaneous activity and blood glucose in LPS-treated mice, 2-DG on its own did not significantly change locomotor activity (Figure 2J) or blood glucose concentrations (Figure 2K) and, on subjective examination, mice receiving 2-DG alone did not appear sick. We have thus demonstrated that reduced glucose metabolism is a key proximate cause of LPS-induced suppression of spontaneous activity.

### Neurodegeneration increases susceptibility to cognitive impairments due to reduced glucose availability

We have previously shown, using the ME7 model, that evolving neurodegeneration progressively increases susceptibility to LPS-induced transient working memory impairments on a T-maze task (Murray et al., 2012; Skelly et al., 2018). We replicate this here to illustrate the time course of these changes, and confirm that LPS does not produce such deficits in normal animals (Figure 3A). We hypothesized that this cognitive vulnerability in ME7 mice may be explained by a greater tendency towards metabolic insufficiency and that cognitive function in ME7 mice might be less able to cope with limiting glucose. We tested this hypothesis using insulin (11.5 IU/kg; i.p.), which significantly lowered blood glucose in both ME7 and NBH mice (Figure 3B). Basal levels of insulin were equivalent in ME7 and NBH mice, and all mice showed similar insulin pharmacokinetics upon insulin treatment (Figure 3C). Despite this, and analogous to LPS-induced cognitive deficits (Figure 3A), insulin induced significant acute working memory dysfunction in ME7 mice that was absent in NBH controls (Figure 3D).

**Figure 3:**
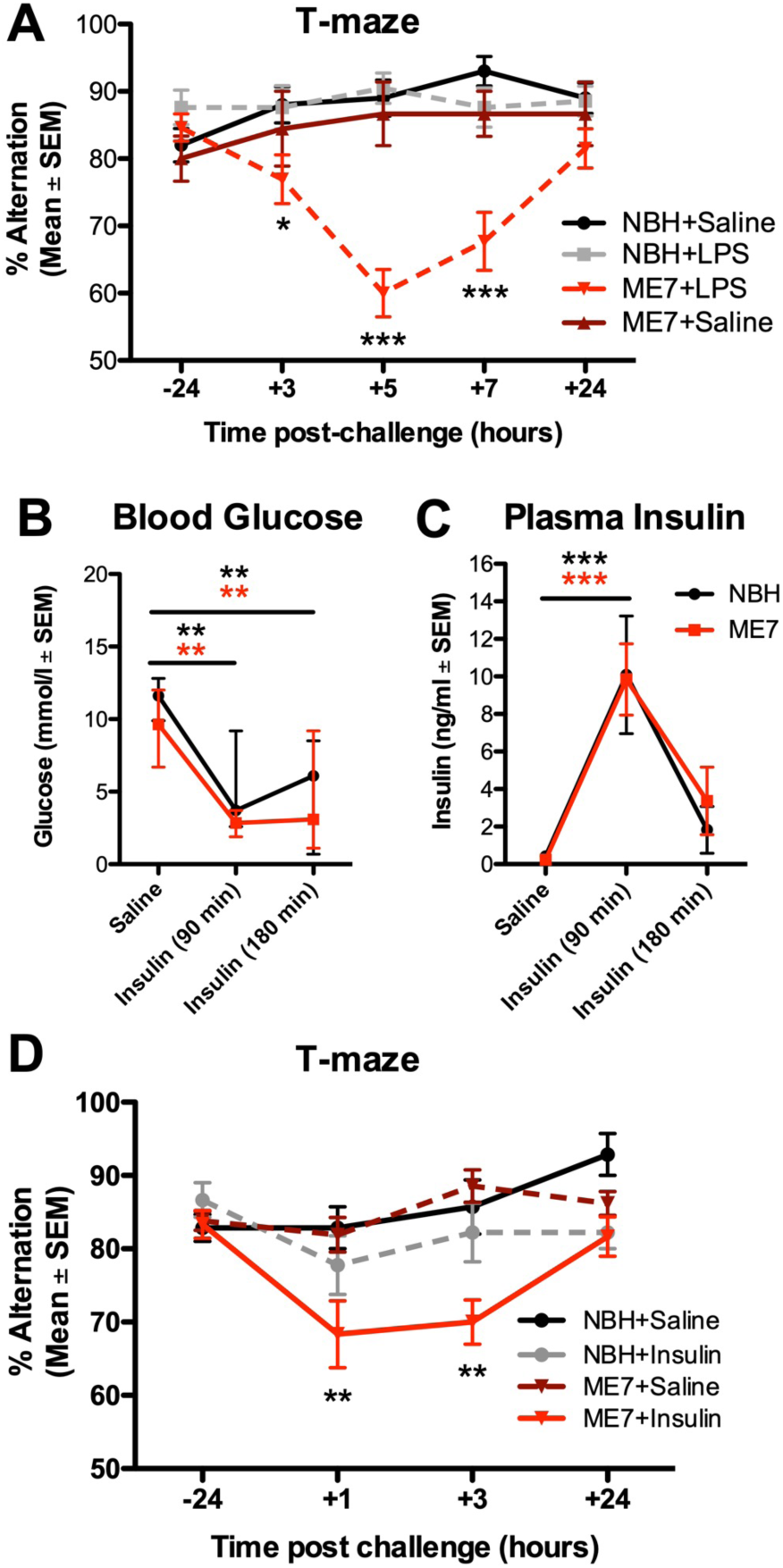
Insulin-induced reductions in blood glucose produces acute cognitive dysfunction selectively in mice with prior neurodegeneration. **A:** ME7 mice have a cognitive vulnerability under LPS treatment (n = 26) that was not present in NBH mice treated with LPS (n = 21). Saline does not induce cognitive deficits either in NBH (n = 20) or in ME7 mice (n = 9). There was an interaction between treatment group and time (*F*_*(12,288)*_ = 5.00; *p* < 0.0001). **B:** Blood glucose (mmol/l) and **C:** Plasma insulin concentrations in saline- or Insulin-treated (11.5 IU/kg; i.p.) NBH and ME7 mice. There were similar reductions in blood glucose (B: main effect of insulin, *F*_*(2,20)*_ = 17.11; *p* < 0.0001) and equivalent insulin concentrations over 180 min in ME7 and NBH animals (C: main effect of insulin, *F*_*(2,28)*_ = 22.86; *p* < 0.0001). **D:** T-maze alternation in ME7 and NBH mice post-challenge with saline or insulin (+1 h = 40-160 min; and +3 h = 160-300 min post-insulin). Testing was performed earlier than in LPS-treated mice as insulin produces a more rapid decrease in blood glucose. There was a significant main effect of insulin (*F*_*(3,135)*_ = 7.418; *p* = 0.0004) and an interaction of ME7 and insulin (*F*_*(9,135)*_ = 3.050; *p* = 0.0024). ME7+insulin-treated mice (n = 12) had significantly lower alternation scores compared to NBH+saline controls (n = 7) at 1 and 3 h post-injection (NBH+insulin: n = 9; ME7+saline: n = 13).

Given the ability of insulin-induced hypoglycemia to trigger cognitive deficits selectively in mice with existing neurodegenerative disease (ME7), we examined whether LPS produced differential hypoglycemic responses in NBH and ME7 animals. Mice were inoculated with ME7 or NBH and, 16 weeks later, challenged with saline or LPS (100 μg/kg; i.p.). LPS produced similar glucose reductions in NBH and ME7 mice in both blood (Figure 4A) and in CSF (Figure 4B), although baseline CSF glucose concentration was slightly higher in ME7 animals with respect to NBH. CSF lactate levels were similar in all 4 groups (Figure 4C). Since ME7 and NBH mice showed equivalent reduction in glucose, but differential cognitive outcomes post-LPS (Figure 3A), and because LPS-induced sickness behavior can be reversed by i.p. glucose (Figure 2E), we hypothesized that the LPS-induced cognitive impairment in ME7 mice might be mediated by limiting glucose supply/utilization. ME7 mice were trained on the ‘escape from water’ T-maze, until criterion performance of >80 % correct was achieved. They were then treated with saline or LPS (100 μg/kg; i.p.) and, 2.5 hours after LPS, treated with saline or glucose (2 g/kg; i.p.) before undergoing T-maze testing. Neither saline-nor glucose-treated ME7 mice deviated from baseline T-maze performance in the absence of LPS, but LPS-treated ME7 mice showed robust impairment between 3-7 hours post-LPS. Those impairments in ME7+LPS+saline mice were significantly attenuated by glucose applied 2.5 hours after LPS (Figure 4D; significant interaction of LPS and glucose: *F*_*(12,240)*_=3.740; *p*<0.0001). Bonferroni *post-hoc* analysis showed that ME7+LPS+glucose mice were significantly different from ME7+LPS+Saline mice at 5 hours.

**Figure 4:**
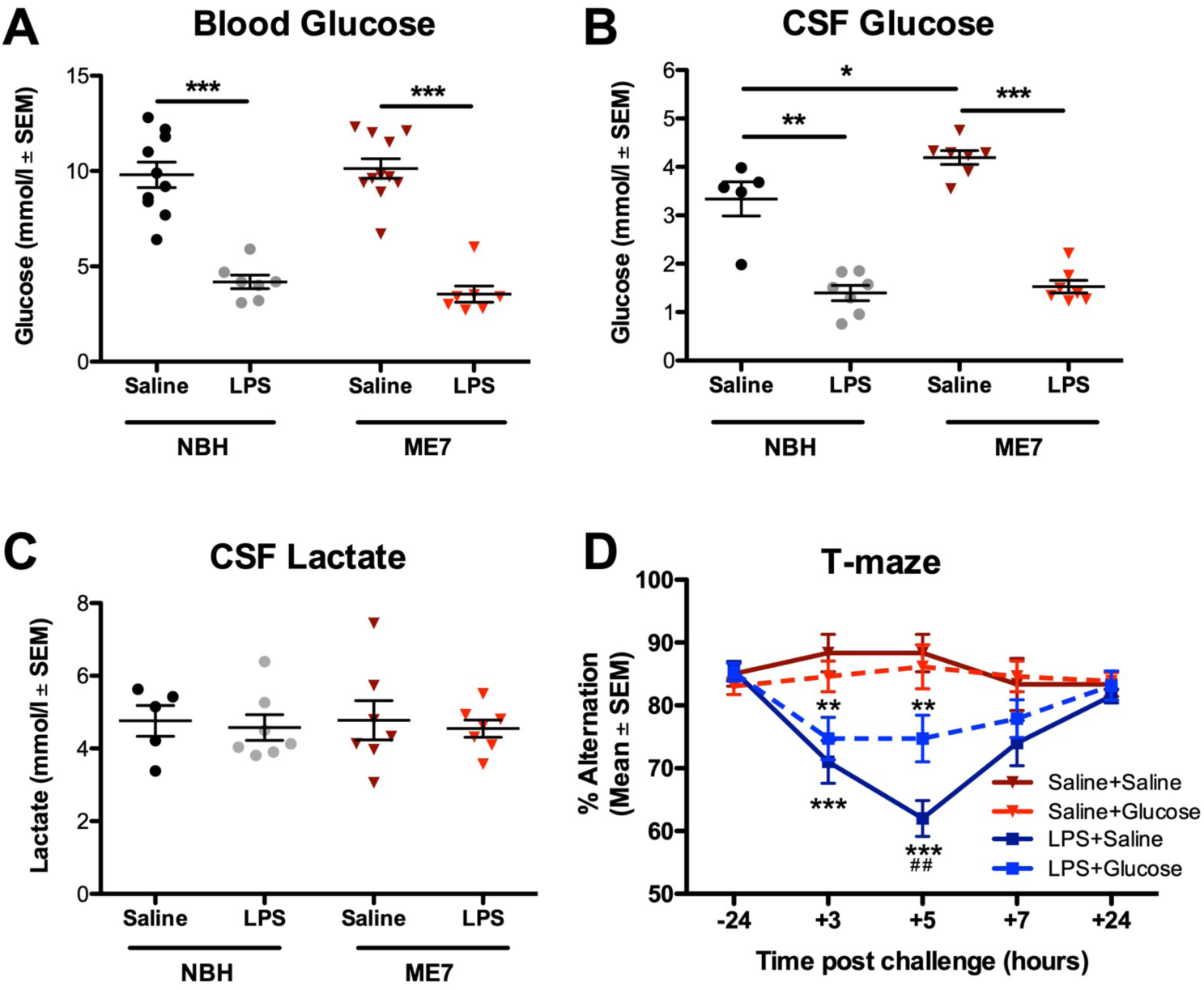
LPS-induced cognitive dysfunction in mice with prior neurodegeneration can be ameliorated by glucose administration. **A:** After 5 hours, LPS produced equivalent decrease in glucose concentration in blood and **B:** CSF glucose in ME7 (n = 7) and NBH mice (n = 7) compared to their respective saline-treated controls (n = 11 and n = 10 respectively). There were main effects of LPS on blood glucose (*F*_*(1,31)*_ = 118.3; *p* < 0.0001) and on CSF glucose (*F*_*(1,22)*_ = 146.5; *p* < 0.0001) and also an effect of disease on CSF glucose (*F*_*(1,22)*_ = 6.665; *p* = 0.0170), with ME7+Saline > NBH+Saline by *post hoc* analysis. **C:** There were no differences in CSF lactate levels. **D:** T-maze alternation in ME7 mice post-challenge with saline or LPS, co-treated with glucose (2 g/kg) or saline. LPS+Saline group (n = 20) showed robust cognitive impairment but the LPS+Glucose group (n = 19) showed significant attenuation. Two-way repeated measures ANOVA showed a main effect of LPS (*F*_*(3,240)*_ = 13.75; *p* < 0.0001) and an interaction of LPS and glucose (*F*_*(12,240)*_ = 3.740; *p* < 0.0001). LPS+Glucose mice performed significantly better than the LPS+Saline group at 5 h post-challenge (^##^*p* < 0.01). Significance levels for Bonferroni *post hoc* tests: \**p* < 0.05; \*\**p* < 0.01; \*\*\**p* < 0.001.

### Human delirium triggered by acute inflammatory trauma (hip fracture) is associated with altered carbohydrate metabolism

Thus, acute inflammation disrupted glucose metabolism and this was causal in acute cognitive dysfunction (Figure 4). We have previously demonstrated that this LPS-induced cognitive deficit is acute, transient and fluctuating, occurs only in animals with prior degenerative pathology and represents the best validated animal model of delirium superimposed on dementia (Davis et al., 2015; Schreuder et al., 2017). Therefore, seeking to investigate generalizability of these findings from mice, we assessed CSF concentrations of glycolytic metabolites in a cohort of acute hip-fracture patients admitted for hip fracture repair with spinal anesthesia (see Table 1 for patient information). This represents an ideal cohort because CSF sampling is possible at the time of spinal anesthesia and because delirium occurs in a significant subset of these patients, and an acute inflammatory trauma (fracture) has been the proximate trigger for this delirium (Hall et al., 2018).

**Table 1.** Demographic information for patients recruited to the study.

|  | No Delirium<br>(n = 32) | Delirium<br>(n = 40) | p-value<sup>d</sup> |
| --- | --- | --- | --- |
| Median age, years (range) | 84.5 (60-93) | 85 (68-95) | 0.69 |
| Male, n (%) | 8 (25) | 11 (27.5) | 0.81 |
| Dementia, n (%)<sup>a</sup> | 2 (6.3) | 32 (80) | <0.001 |
| Independent in activities of daily living, n (%)<sup>b</sup> | 23 (71.9) | 8 (20) | <0.001 |
| Living in an institution, n (%) | 3 (9.4) | 20 (50) | <0.001 |
| APACHE II, median (IQR)<sup>c</sup> | 8 (6.3 – 9.8) | 9 (8 – 11) | 0.004 |
| CCI, median (IQR) | 1 (0 – 1.8) | 1 (0 – 2) | 0.044 |
| ASA score, median (IQR) | 2 (2-3) | 3 (3-3) | <0.001 |

<sup>a</sup>Based upon consensus in an expert panel. <sup>b</sup>Defined as 19 or 20 points on Barthel Activities of Daily Living. <sup>c</sup>Without information on haematocrit and arterial blood gas. <sup>d</sup>Mann-Whitney test and Chi-square tests depending on data distribution. APACHE II = Acute Physiology and Chronic Health Evaluation II; ASA = American Society of Anesthesiologists Physical Health Classification; CCI = Charlson Comorbidity Index score; IQR = Interquartile Range.

Hip-fracture patients were assessed for delirium at the time of lumbar puncture and those with prevalent delirium were compared to those without delirium on CSF glucose, lactate and pyruvate (commonly used markers of central energy metabolism disturbance in clinical populations (Leen et al., 2012; Zhang and Natowicz, 2013)). CSF glucose was not different in those with and without delirium (Figure 5A). A previous study of all-cause delirium versus stable dementia (Caplan et al., 2010) provided the *a priori* hypothesis that delirium would be associated with elevated lactate, and lactate was indeed significantly elevated during delirium (Figure 5B; one-tailed Mann-Whitney analysis; *p* = 0.0128). Changes in CSF lactate, associated with delirium, were not explained by dementia status (data not shown). Median pyruvate levels were significantly elevated in delirium (Figure 5C) and although pyruvate was not detectable in CSF of all patients, it was detected significantly more often in patients with delirium (data not shown). Increases in the CSF lactate:glucose ratio (LGR) have been associated with reduced consciousness (Sanchez et al., 2013) and increased mortality (Lozano et al., 2019) and here these data indicate an elevated LGR both in humans experiencing delirium after acute inflammatory trauma (Figure 5D) and in mice cognitively impaired by acute systemic inflammation (Figure 5E). The changes in LGR observed in mice and humans differ in how they arise, with an increase in the ratio driven by increases in lactate in humans (Figures 5A and 5B) and by decreases in glucose in the mouse (Figure 4A and 4C), but both mouse and human data sets indicate that there is a significant derangement of brain energy metabolism following these inflammatory insults and, in mice, this is clearly causal for acute cognitive dysfunction.

**Figure 5:**
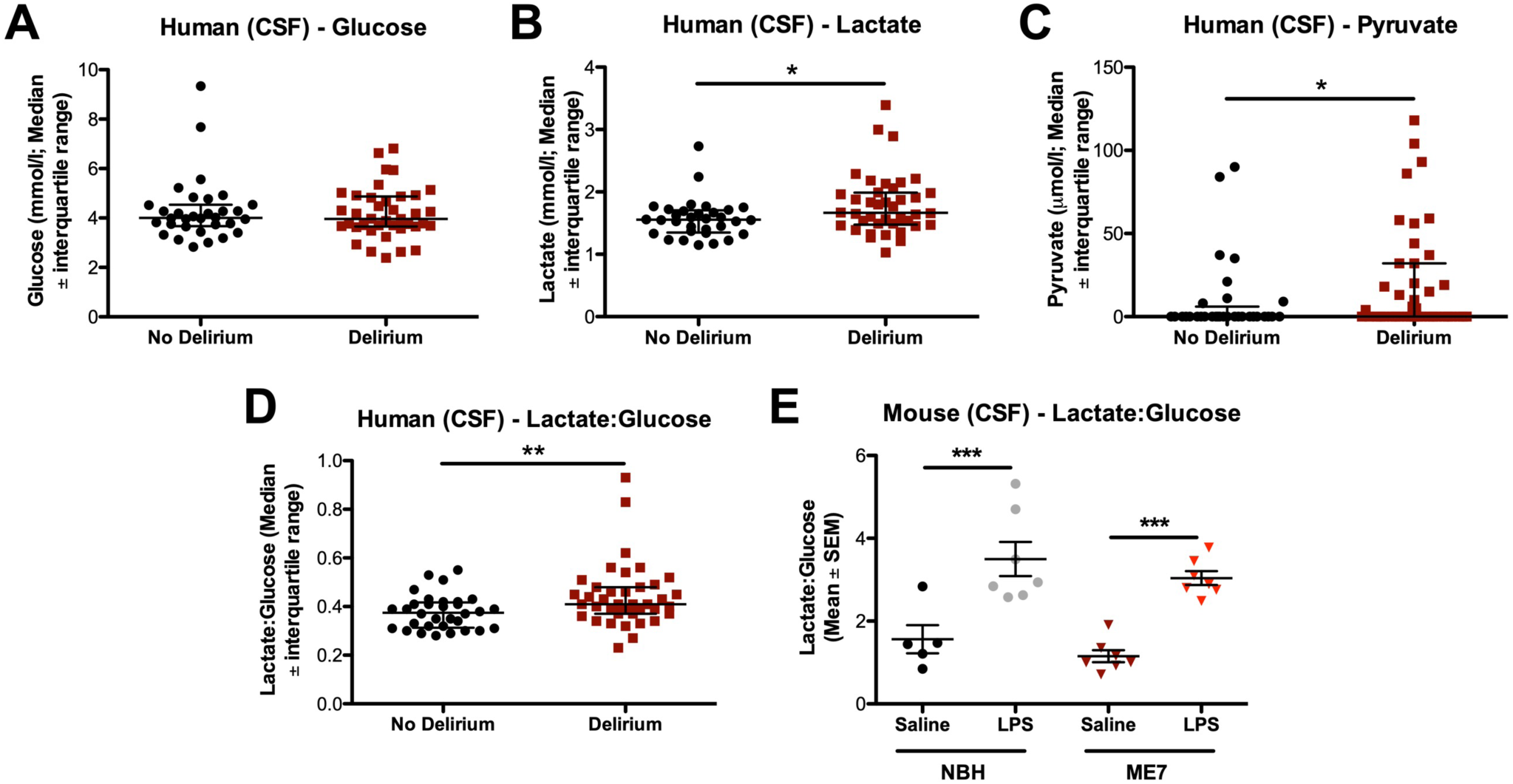
Derangement of energy metabolism in human delirium. Metabolite levels in the CSF of hip fracture patients with delirium (n = 40) at the time of CSF sampling compared to age-matched patients with no delirium at any point of their hospital stay (n = 32). **A:** Glucose levels in delirium (n = 39, 1 sample omitted due to a read error) and non-delirium cases were not significantly different (Mann-Whitney *U* = 606.5; *p* = 0.8442). **B:** Patients with delirium had significantly higher levels of lactate in their CSF compared to controls (*U* = 442.5; *p* = 0.0128). **C:** Patients with delirium showed significantly higher pyruvate levels compared to controls (*U* = 514.5; *p* = 0.0494). **D:** The lactate:glucose ratio (LGR) for patients with delirium (n = 39) was significantly higher compared to controls (*U* = 399.5; *p* = 0.0048). **E:** LPS significantly increased the CSF LGR in both ME7 (n = 7) and NBH mice (n = 7) compared to their respective saline-treated controls (n = 7 and n = 5; *F*_(1,22)_ = 44.58; *p* < 0.0001). Significance levels for Bonferroni *post hoc* tests: \*\**p* < 0.01; \*\*\**p* < 0.001.

## Discussion

We demonstrate that LPS-induced hypoglycemia suppresses spontaneous activity in mice. Glycemic status was a major determinant of spontaneous activity after LPS. Reduced glucose availability also drove LPS-induced acute cognitive impairment in mice with underlying neurodegeneration. The degenerating brain is also selectively vulnerable to cognitive disruption by insulin, despite equivalent blood glucose reductions. Finally, inflammatory trauma-induced delirium in humans was associated with altered central energy metabolism.

### Hypoactivity

Reducing blood glucose can be adaptive for the organism, depriving infectious agents of a key fuel source, and providing further glucose can actually increase *Listeria monocytogenes*-induced mortality (Wang et al., 2016). Nonetheless, we show, in the acute phase, that significant LPS-induced decreases in blood glucose reduce CSF glucose and suppress spontaneous activity. This is consistent with prior studies showing correlation between blood glucose and sickness behavior (Carlton and Demas, 2017) and others showing that insulin-induced hypoglycemia supresses social activity in c57BL/6 mice (Park et al., 2008; Park et al., 2012). Here, by directly increasing glucose availability, we prevent LPS-induced suppression of activity without reducing IL-1β. Moreover, 2-DG completely blocked LPS-induced IL-1β secretion, as previously shown for macrophage IL-1 production (Tannahill et al., 2013), but in also preventing glucose utilization it further suppressed activity. Although IL-1 is widely implicated in LPS-induced sickness behavior, LPS-induced hypoactivity persisted even when IL-1 secretion or action was blocked (IL-1RA, IL-1RI^-/-^). Although IL-1RA protection previously inhibited LPS-induced hypoglycemia, those effects were partial: 100 μg IL-1RA/mouse marginally mitigated hypoglycemia (Del Rey et al., 2006), as we observed here. IL-1RA (at 200 μg/mouse) completely blocked effects of 25 μg/kg IL-1β (Figure 1D), which leads to blood levels of approximately 600 pg/ml IL-1β (Skelly et al., 2013), and this IL-1RA dose should therefore block the effects of LPS-induced IL-1β (∼50-100 pg/ml) (Teeling et al., 2007; Murray et al., 2011; Skelly et al., 2013). However, TNFα also triggers hypoglycemia (Oguri et al., 2002) thus IL-1RA can have only a partial effect in limiting LPS-induced hypoglycemia. Therefore, while IL-1β might be a key mediator of hypoglycemia at 25 μg/kg LPS (Del Rey et al., 2006), IL-1RA barely limits hypoglycemia with LPS at 250 μg/kg (current study). Ultimately the ability of glucose to restore activity in the current experiments reveals the importance of glucose uptake and use in fueling and regulating spontaneous activity under LPS-induced inflammation.

The hypothalamus, monitors levels of circulating IL-1β (Matsuwaki et al., 2017) and glucose (Lopez-Gambero et al., 2018) and coordinates sickness behavior. IL-1β action in the hypothalamus is proposed to reprogram the organism to operate at lower circulating glucose levels after LPS (25 μg/kg) (Del Rey et al., 2006) and these authors propose that IL-1β increases brain energy metabolism (Del Rey et al., 2016). [^18^F]-fluorodeoxyglucose (FDG-PET) experiments show that high dose LPS (15 mg/kg) increased hypothalamic activity (Wang et al., 2016) but LPS (10 mg/kg i.p.) also decreased glucose uptake across multiple cortical regions (Semmler et al., 2008). If, as reported, IL-1 lowers the set-point for glucose homeostasis, allowing animals to function efficiently at lower glucose concentrations (Del Rey et al., 2006), it is not intuitive why transiently increasing available glucose should rapidly increase spontaneous activity. However, here we show that administration of glucose raises both blood and CSF glucose (Figure 2). Although this “top-up” of glucose provides only temporary and partial increases in available glucose (Figure 2G) (Del Rey et al., 2006), this is sufficient to restore spontaneous activity and cognition. We therefore propose that while the hypothalamus might be selectively active during acute inflammation, to coordinate neuroendocrine responses to the acute threat, the suppression of spontaneous locomotor activity that is actually observed may reflect decreased neural activity underpinned by decreased available glucose.

The neuroanatomical basis of LPS-induced suppression of exploratory activity is incompletely understood but correlates with suppression in cFOS in brain areas associated with positive motivation (Stone et al., 2006) and exploratory behavior (Gaykema and Goehler, 2011). LPS triggers norepinephrine (NE) release in the hypothalamus (Francis et al., 2001) and lesioning caudal medullary NE inputs to the hypothalamus blocks LPS-induced hypoactivity (Gaykema and Goehler, 2011). Hypoglycemia, hyperinsulinemia (Beverly et al., 2001) and 2-DG treatment (Beverly et al., 2000) all induce hypothalamic NE release, suggesting potential points of convergence for how inflammation and impaired glucose metabolism may drive changes in behavior during sickness.

Whatever the neuroanatomical and neurotransmitter underpinnings, the current data strongly support the idea that available and usable glucose is a key determinant of LPS-induced suppression of activity. This has implications for studies using peripheral LPS to examine the neurophysiological and behavioral consequences of systemic infection. Levels of circulating LPS arising from bolus LPS challenges are higher than in active infection (Danner et al., 1991). Bolus LPS treatment (4 ng/kg; i.v.) in human volunteers transiently decreases plasma glucose (Bloesch et al., 1993) while active infection typically does not produce hypoglycemia (Furman et al., 1988). Therefore, although bolus LPS would appear to have face validity as a model of systemic infection, if key behavioral and neurophysiological changes induced by LPS in experimental subjects are underpinned by a physiological change, i.e. hypoglycemia, that rarely occurs during active infection, this necessitates a review of the generalizability of bolus LPS-induced changes to understand changes during active infection.

### Acute cognitive dysfunction and delirium

Human data suggest that reduced glucose uptake in the medial temporal lobe associates with impaired performance in hippocampal-dependent tasks (Harrison et al., 2014). Remarkably, despite the robust and long lasting reductions in available glucose shown here, normal LPS-treated mice maintain good working memory (Skelly et al., 2018). The same decreases in glucose, caused by LPS or insulin, were sufficient to trigger dysfunction in animals with prior neurodegeneration. Speculatively, the circuitry underpinning working memory may, during neurodegeneration, operate close to thresholds for decompensation and this task may then need to recruit additional brain areas to maintain sufficient working memory. Exogenously added glucose does not improve cognition in young rats (Kealy et al., 2017) but enhances cognition in aged rats (McNay and Gold, 2001), supporting the idea that the same task may require additional metabolic support in the aged or degenerating brain. The addition of a further stressor may be sufficient to unmask underlying vulnerability. Volunteers exposed to *Salmonella Typhi*. vaccination performed equally to controls on the Stroop test of executive function but recruited additional areas of the prefrontal and anterior cingulate cortex to maintain performance during inflammation (Harrison et al., 2009). If increased connectivity is required to maintain performance during inflammation, then inflammatory insults may unmask vulnerability, when evolving neurodegeneration impairs connectivity (Davis et al., 2015). Until now, the ME7 model of delirium during dementia has been an exemplar for an inflammatory hypersensitivity, but these data show that they are also more vulnerable to bioenergetic stressors. Despite equivalent reductions in blood and CSF glucose, NBH animals are resilient to hypoglycemia-induced cognitive impairment but ME7 animals are vulnerable, whether induced by LPS or by insulin.

The brain is a metabolically demanding organ and it may be adaptive, for survival, to minimize energy use in the brain and preserve autonomic function at the expense of higher cortical function. Engel and Romano proposed that delirium is driven by a failure to meet the brain’s energy requirements, regardless of the underlying cause (Engel and Romano, 2004). Hypoglycemia is sufficient, alone, to produce delirium and EEG slowing and this is reversed by glucose administration (Engel and Romano, 1944). Small CSF studies support the idea of metabolic disturbances during delirium: patients with delirium have elevated CSF lactate compared to non-delirious Alzheimer’s disease controls (Caplan et al., 2010) and [^18^F]FDG-PET studies show decreased glucose uptake (Haggstrom et al., 2017). The posterior cingulate cortex, which is associated with attention and arousal, was particularly affected and disrupting energy metabolism here could be important in delirium.

Although hypoglycemia in mice is not precisely defined, the ME7 blood glucose concentrations here remained just above the clinical threshold for moderate hypoglycemia (3.9 mmol/l) (Cryer, 2017). Iatrogenic hypoglycemia, is common in patients using insulin for diabetes (Cryer, 2002) and is a major cause of emergency department admissions and adverse CNS effects in older patients (Shehab et al., 2016). Significantly, our ME7 data suggest that even when blood glucose levels do not fall into classical hypoglycemic ranges, these changes have significant deleterious impacts on brain function in those with prior vulnerability.

It is important to note that CSF glucose was not lower in the hip fracture patients studied here (Figure 5A). Although both hypoglycemia and hyperglycemia strongly increase risk for sepsis-associated encephalopathy (Sonneville et al., 2017), hypoglycemia is less common. However, hyperglycemia-associated delirium typically occurs during insulin insensitivity, where glucose uptake and use is impaired. In humans, insulin resistance occurs upon LPS (Agwunobi et al., 2000), infection (Virkamaki et al., 1992) and surgery (Thorell et al., 1994). Microcirculatory failure and tissue hypoxia, which commonly occur in sepsis (Ince and Mik, 2016) may also drive inefficient glucose oxidation. The higher levels of lactate and pyruvate in the current study may indicate a shift from normal aerobic to anaerobic glycolysis. Although dementia status is a major risk factor for delirium (Davis et al., 2015), the changes in lactate and LGR observed are not explained by the presence of dementia in these patients. The data are consistent with previously demonstrated associations between delirium and increased CSF lactate (Caplan et al., 2010) and hypoxia (Tahir et al., 2018).

### Conclusion

Reduced glucose availability is a major driver of LPS-induced suppression of spontaneous activity. In animals made vulnerable by evolving neurodegeneration, this decreased glucose is now sufficient to trigger acute cognitive dysfunction indicating that metabolic insufficiency underlies cognitive dysfunction in this animal model resembling delirium. A disruption of energy metabolism also occurs in delirium triggered by inflammatory trauma. Together, the findings indicate that disrupted energy metabolism contributes to general behavioral changes associated with sickness but also to acute neuropsychiatric disorders such as delirium. These data should focus attention on bioenergetic mechanisms of acute brain failure during acute illness and hospitalization in older adults.

## Acknowledgements

This work was supported by The Wellcome Trust (CC SRF 090907) and by the US National Institutes of Health (R01AG050626). JPL acknowledges financial support from the Health Research Board (HRB EQ/2004/12). LOW is funded by the Norwegian Health Association and by the South-Eastern Norway Regional Health Authorities. Thank you to Prof. Stuart Allan for the gift of IL-1RA and Prof. Kingston Mills for the gift of IL-1RI^-/-^ mice. This article was made available as a preprint on BioRxiv (https://doi.org/10.1101/642967).

